# Loss-of-function in *IRF2BPL* is associated with neurological phenotypes

**DOI:** 10.1101/322495

**Authors:** Paul C. Marcogliese, Vandana Shashi, Rebecca C. Spillmann, Nicholas Stong, Jill A. Rosenfeld, Mary Kay Koenig, Julián A. Martínez-Agosto, Matthew Herzog, Agnes H. Chen, Patricia I. Dickson, Henry J. Lin, Moin U. Vera, Noriko Salamon, Damara Ortiz, Elena Infante, Wouter Steyaert, Bart Dermaut, Bruce Poppe, Hyung-Lok Chung, Zhongyuan Zuo, Pei-Tseng Lee, Oguz Kanca, Fan Xia, Yaping Yang, Edward C. Smith, Joan Jasien, Sujay Kansagra, Gail Spiridigliozzi, Mays El-Dairi, Robert Lark, Kacie Riley, Dwight D. Koeberl, Katie Golden-Grant, Program for Undiagnosed Diseases (UD-PrOZA) Undiagnosed Diseases Network, Shinya Yamamoto, Michael F. Wangler, Ghayda Mirzaa, Dimitri Hemelsoet, Brendan Lee, Stanley F. Nelson, David B. Goldstein, Hugo J. Bellen, Loren D.M. Pena

## Abstract

The *Interferon Regulatory Factor 2 Binding Protein Like* (*IRF2BPL*) gene encodes a member of the IRF2BP family of transcriptional regulators. Currently the biological function of this gene is obscure, and the gene has not been associated with a Mendelian disease. Here we describe seven individuals affected with neurological symptoms who carry damaging heterozygous variants in *IRF2BPL.* Five cases carrying nonsense variants in *IRF2BPL* resulting in a premature stop codon display severe neurodevelopmental regression, hypotonia, progressive ataxia, seizures, and a lack of coordination. Two additional individuals, both with missense variants, display global developmental delay and seizures and a relatively milder phenotype than those with nonsense alleles. The bioinformatics signature for *IRF2BPL* based on population genomics is consistent with a gene that is intolerant to variation. We show that the *IRF2BPL* ortholog in the fruit fly, called *pits* (*protein interacting with Ttk69 and Sin3A*), is broadly expressed including the nervous system. Complete loss of *pits* is lethal early in development, whereas partial knock-down with RNA interference in neurons leads to neurodegeneration, revealing requirement for this gene in proper neuronal function and maintenance. The nonsense variants in *IRF2BPL* identified in patients behave as severe loss-of-function alleles in this model organism, while ectopic expression of the missense variants leads to a range of phenotypes. Taken together, *IRF2BPL* and *pits* are required in the nervous system in humans and flies, and their loss leads to a range of neurological phenotypes in both species.

## Introduction

The etiology of neurodevelopmental disorders can vary and can include prenatal exposures, maternal disease, multifactorial causes and single genes. *De novo* genomic contributions were recently highlighted in a large cohort studied as part of the Deciphering Developmental Disorders Study, with an estimate that 42% of the cohort carried pathogenic *de novo* mutations in the coding region of genes.^1-3^ The phenotypic and genomic heterogeneity of neurodevelopmental disorders can pose a diagnostic challenge. Several known Mendelian disorders such as Rett syndrome (MIM: 312750), the neuronal ceroid lipofuscinoses (NCLs), and X-linked adrenoleukodystrophy (X-ALD, MIM: 300100) display neurodevelopmental regression as a common element and illustrate the range of symptoms and pathologies associated with neurological symptoms. The known or proposed mechanisms of disease can include altered transcriptional control (Rett syndrome)^4,5^, accumulation of a substrate with loss of neurons (juvenile infantile NCL, MIM: 204200)^6,7^, or inflammation and demyelination (X-ALD).^8-10^ More recently, new genes associated with severe developmental phenotypes and neurodegeneration have been discovered.^11,12^ This has been largely possible due to the use of high-throughput sequencing methods such as next-generation sequencing (NGS), in conjunction with sequencing databases for control cohorts such as ExAC and gnomAD^13-21^, variant prediction^22^, model organism information (e.g. MARRVEL.org)^23^ and crowdsourcing programs to identify additional cases, such as GeneMatcher.org.^24^ These tools have greatly promoted gene discovery and assisted in ascertaining the role of the candidate variants for disease. Programs such as the Undiagnosed Diseases Network (UDN)^25,26^, promote multi-site collaboration that combines NGS and functional data that facilitate the diagnosis of rare disorders.

Here we describe a cohort of seven cases, ascertained for neurological symptoms, who share predicted pathogenic variants in *IRF2BPL*, an intronless gene at 14q24.23.^27^ The transcript is expressed in many organs, including in the central nervous system (CNS) components such as the cerebellum (GTex, accessed 1/29/2018).^28^ The IRF2BPL protein and its two mammalian paralogs, IRF2BP1 and IRF2BP2, share two highly conserved domains. These include a coiled-coil DNA binding domain (IRF2BP zinc finger domain) at the amino-terminus and a C3HC4-type RING finger domain at the carboxy-terminus. IRF2BPL also contains polyglutamine and polyalanine tracts. In between the two conserved domains is a variable region that contains a potential nuclear targeting signal.^27^ IRF2BPL also contains several putative PEST (proline, glutamic acid, serine, and threonine-rich) sequences throughout the protein suggesting that the IRF2BPL protein is post-translationally regulated^29^ (Figure 1A).

**Figure 1:**
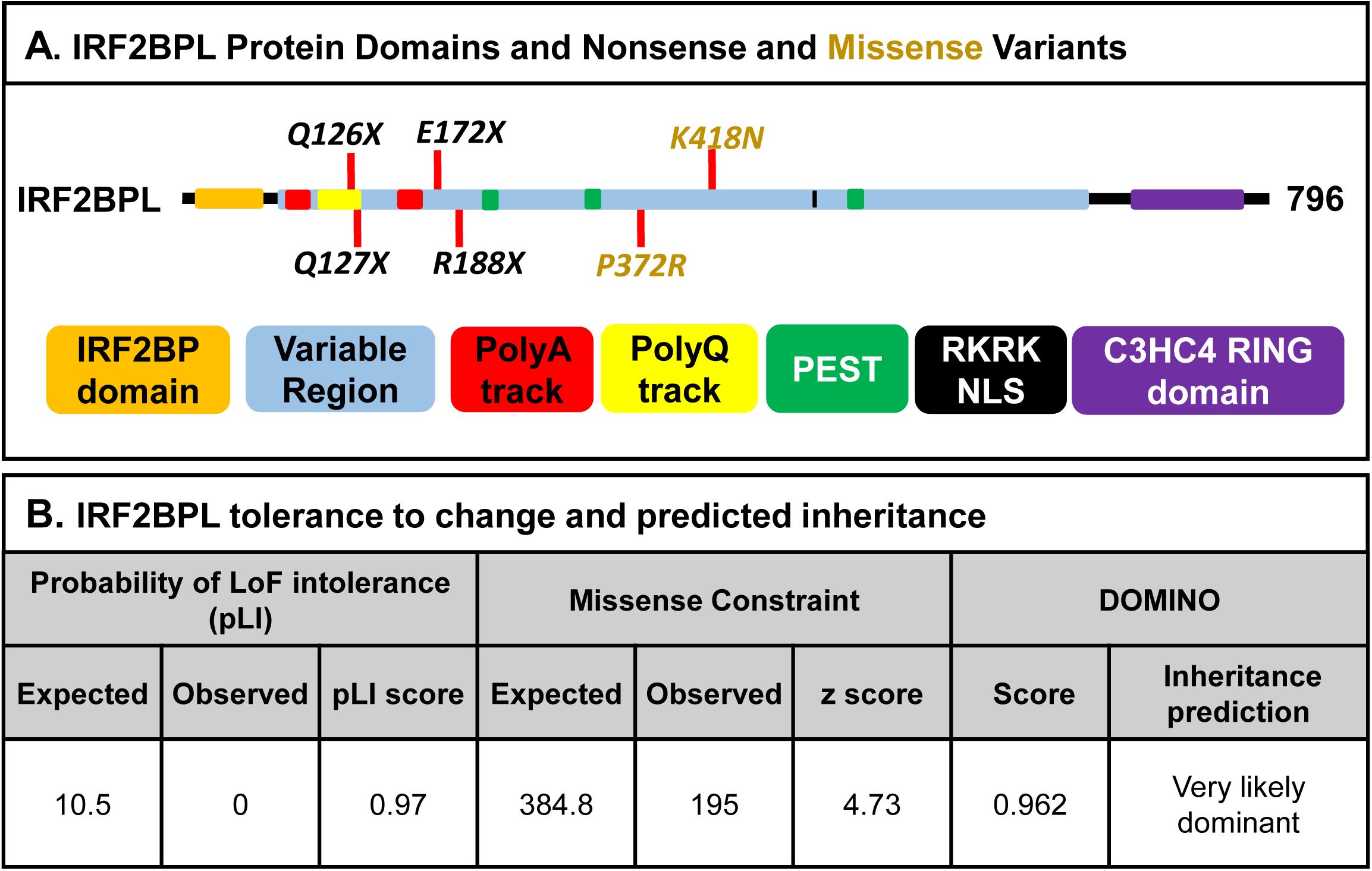
IRF2BPL protein structure and gene constraint. (A) IRF2BPL is 796 amino acids (AA) long and contains two highly conserved domains (IRF2BP zinc finger and C3HC4 RING) and the N- and C-termini. Within the variable region are multiple polyalanine and PEST sequences and a 25 AA polyglutamine tract (AA 103 to 127). All four nonsense variants occur early in the transcript before predicted PEST sequences, and the two missense variants (highlighted in orange) occur in the middle of the protein. (*B*) *IRF2BPL* is highly constrained based on the lack of LoF variants in ExAC ^20^ resulting in a high probability of LoF intolerance (pLI) score and missense constraint z-score. Predictions based on the DOMINO algorithm indicate that variation in IRF2BPL is likely to lead to a dominantly inherited disease.

IRF2BPL has been proposed to have a role in the initiation of puberty in female non-human primates and rodents.^30^ It acts as a transcriptional activator for Gonadotropin Releasing Hormone in the CNS.^30^ Expression of *Irf2bpl* in the hypothalamus of female rats increases during puberty and site-specific reduction of *Irf2bpl* in the preoptic area disrupts the estrus cycle.^31^ In addition, IRF2BPL has also recently been proposed to function as an E3 ubiquitin ligase that targets β-catenin for proteasome degradation in gastric cancer.^32^ Despite these studies, the *in vivo* function of IRF2BPL in all species remains largely undefined. Here we describe a novel role for *IRF2BPL* in the functional and structural maintenance of the nervous system. We provide evidence in seven patients and use functional assays in fruit flies to support the findings that these variants cause dramatic changes to IRF2BPL function and that IRF2BPL plays a role in both development and neuronal maintenance.

## Subjects and Methods

*Demographics, ascertainment, and diagnoses* (*Table 1 and supplementary table 1*). The individuals are all unrelated and between 2.5 and 43 years of age, including one patient who died at 15 years of age. Four are male, six are Caucasian, and one identified as Hispanic. They were evaluated by the genetics or neurology service as part of clinical care for neurological symptoms (subjects 2, 4, 5, 6, 7) or as participants (subjects 1 and 3) in the UDN.^26^ In non-UDN cases, use of the GeneMatcher website^24^ facilitated ascertainment of two of the subjects (6 and 7); the UDN page for *IRF2BPL* facilitated contact with subject 5, and subjects 2 and 4 were ascertained through communication with the clinical lab that performed whole exome sequencing (WES). WES was performed as a trio in five subjects (1, 3, 5, 6, and 7) and proband-only in two (2 and 4). WES was performed with written informed consent for clinical sequencing and in accordance with institutional review board procedures for research sequencing for all subjects. Consent for publication and images were obtained from all guardians.

The seven cases reported here have a constellation of neurological findings of variable severity. While almost all suffered from epilepsy [Human Phenotype Ontology^33^ term (HP: 0001250)], those with nonsense variants in *IRF2BPL* had severe, progressive neurodevelopmental regression (HP: 0002376), dysarthria (HP: 0001260), spasticity (HP: 0001257), and symptoms of movement disorders (HP: 0100022). There was also cerebral volume loss in the two oldest cases (3 and 5; HP: 0002506), and cerebellar volume loss in case 5 (HP:0007360). In contrast, the two cases with missense variants have a generally milder course, with symptoms of epilepsy, speech delay (HP: 0000750), and hypotonia (HP: 0009062). Each proband is described below, along with additional details in Table 1 and Supplementary Table 1.

**Table 1.**
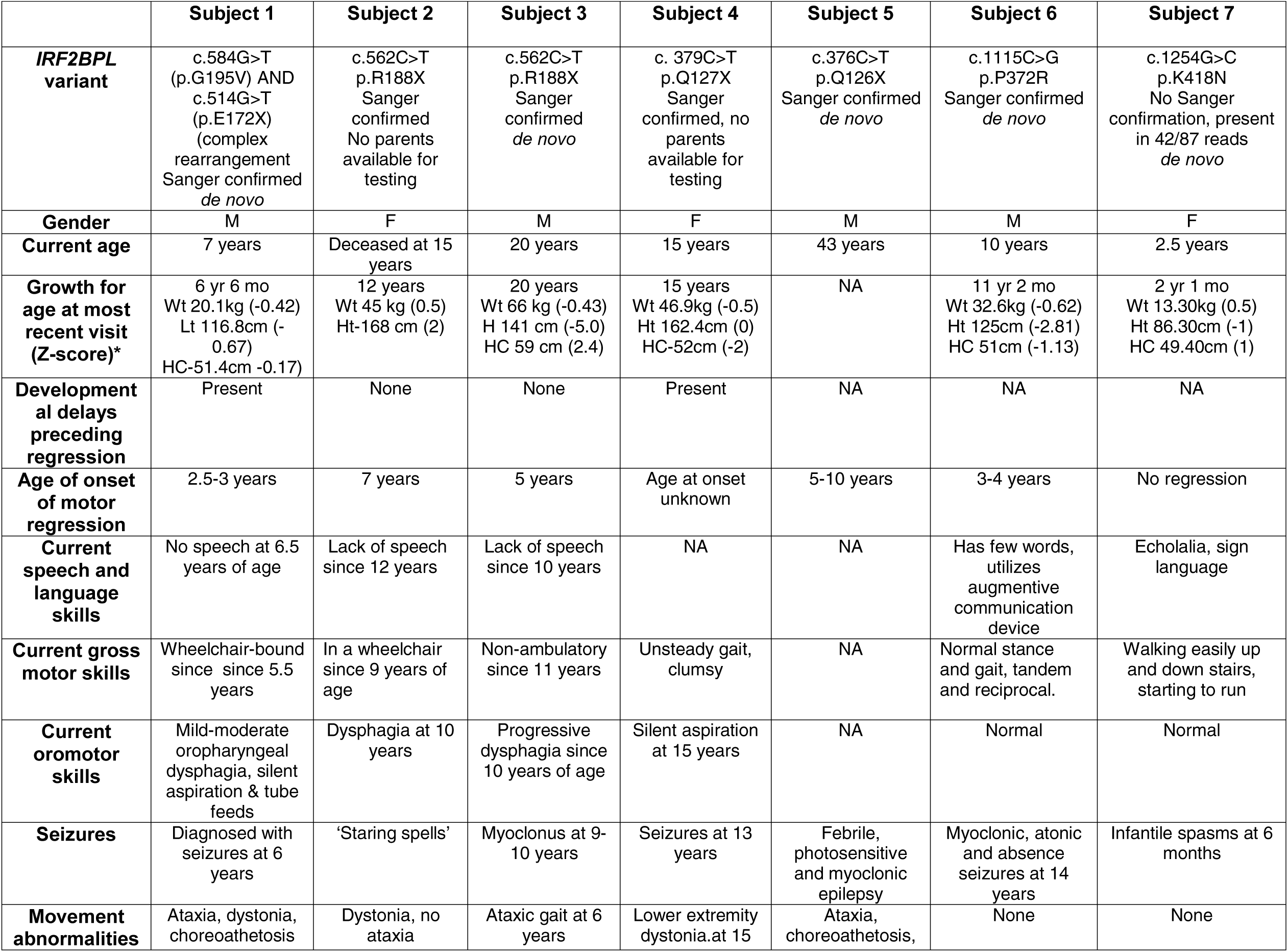
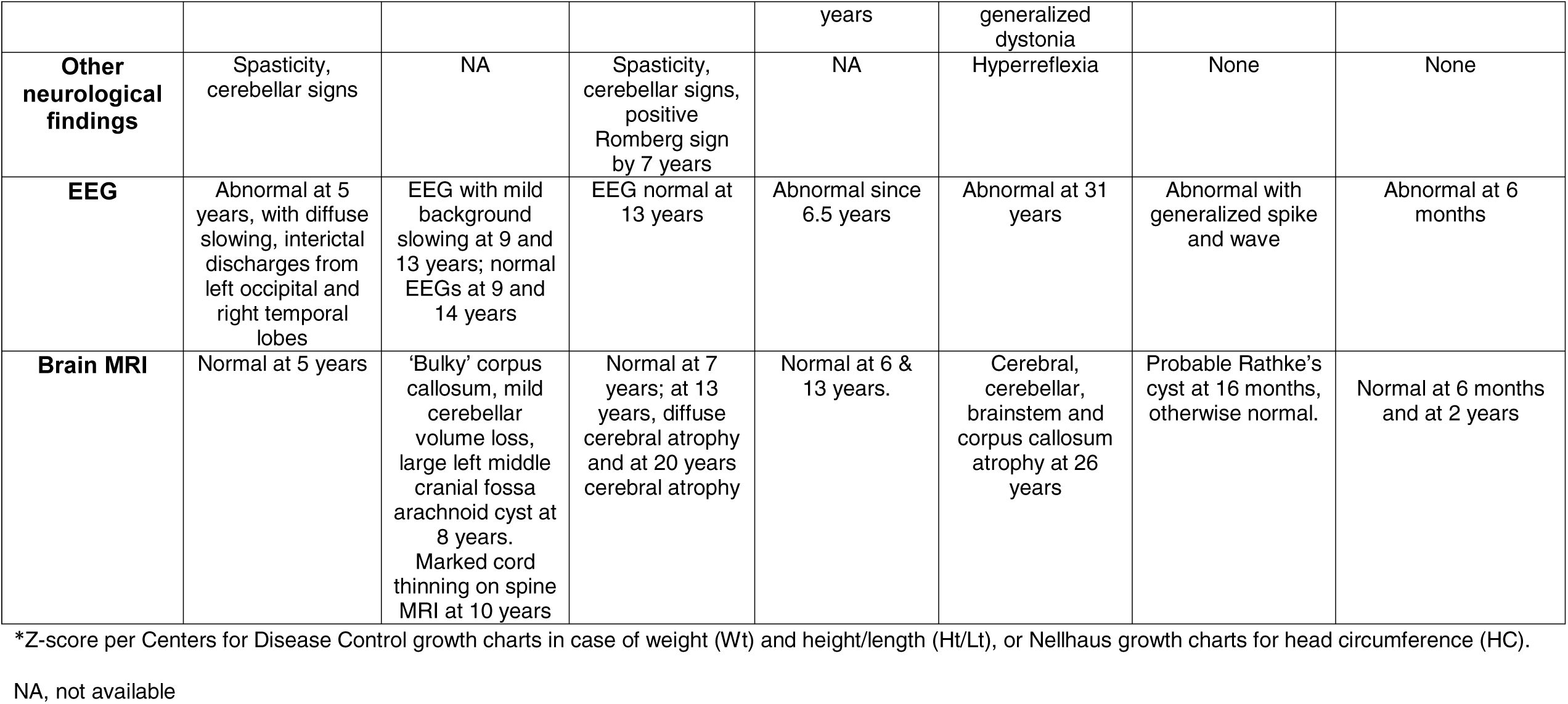
Salient clinical features for each subject.

### Subject #1

This is a 7-year-old male who met early gross and fine developmental milestones in the first two years of life (Supplementary Figure 1). He had a minor speech delay, as he had his first word at 15 months of age. His development was normal until 2.5 years, when clumsiness and excessive falling were noted. At the age of 4 years, the patient was unable to walk without someone holding his hands or using a harness or a walker (Supplemental Movie 1). During an evaluation at 7 years, the patient had lost the ability to walk, was not able to grasp objects, was not able to feed himself and had no expressive language. He had progressive loss of gross and fine motor function, speech and self-help skills. Additionally, he had a diagnosis of dystonia, choreoathetosis, and variable spasticity that primarily affected his lower extremities.

Previous evaluations included a normal brain magnetic resonance imaging (MRI), audiology, and electromyography (EMG) at 5 years of age. He developed esotropia at the age of 4 years that progressively became of a larger angle. Ophthalmic examination at the age of 5 years was notable for visual acuity better than 20/70 in either eye with full fields to distraction. There was evidence of large angle esotropia (higher than 65) and bilateral facial palsies with ophthalmoplegia sparing the eyelids and the pupils. There was a severe horizontal gaze palsy but vertical gaze was also limited. Saccades were decreased in both eyes. Funduscopic examination was normal and he had a normal electroretinogram with normal optical coherence tomography scan of the retina and the retinal nerve fiber layer (RNFL) (measured 85 µm on the RNFL, which is normal for age).

While his neurodevelopmental course demonstrated significant developmental regression, his cognition was tested with the Peabody Picture Vocabulary Test, Fourth Edition, at 6 years of age. He scored in the average range for cognition, consistent with a previous score on the Verbal Ability composite of the Differential Ability Scales-Second Edition, administered when he was 3 years, 9 months of age. Although he did not have clinical seizures at the time, an electroencephalogram (EEG) at 6 years of age showed diffuse slowing that was consistent with generalized brain dysfunction and interictal discharges from the left occipital and right temporal leads, indicating a possible area of epileptogenic potential. At almost 7 years of age, he had an episode of abnormal movements resembling myoclonus and was diagnosed with a seizure disorder, which was controlled with levetiracetam.

Prior biochemical, cytogenetic and molecular evaluations are summarized in Supplementary Table 2. Reanalysis of previously obtained research WES data (trio) at Columbia University identified heterozygosity for a *de novo* truncating variant in *IRF2BPL*, NM_024496.3, c.514G>T (p.E172X) that was Sanger confirmed. Additionally, he had a *de novo* missense variant downstream and in *cis* to the nonsense variant in *IRF2BPL*, NM_024496.3, c.584G>T (p.G195V).

### Subject #2

The patient is a female who met all of her early gross developmental and speech milestones. At 5-6 years of age her family began to notice progressive gait disturbance. Her initial diagnosis by the neurology service was dystonia, and she was given a trial of carbidopa-levodopa (Sinemet) with no improvement. Symptoms progressed, and by 8 years of age she had notable lower extremity spasticity with dysarthria, drooling, dysphagia, and incontinence. Examination at 9 years of age demonstrated an interactive girl with intact cognition, decreased speech fluency and severe dysarthria. Coordination exam demonstrated past pointing. Gait was wide-based with spastic diplegia and scissoring. She had an intrathecal baclofen pump placed at 9 years of age with no improvement. She had lost the ability to stand alone or walk by 10 years and was unable to use her hands or speak by 11 years. She had a dysconjugate gaze at 11 years. MRI spine studies were initially normal but at 10 years demonstrated marked cord thinning. MRI of the brain demonstrated mild cerebellar volume loss at 8 years. The cerebellum was small and the corpus callosum was ‘bulky’. Her respiratory function became compromised and ventilation by continuous positive airway pressure was recommended at 10 years. A G-tube was placed at 10 years for dysphagia and aspiration. During her last formal examination at 13 years of age, her cognition was difficult to assess, as she was no longer able to communicate. All symptoms continued to progress until her family chose to limit her medical interventions for feeding intolerance, and she died at 15 years of age. A nonsense variant in *IRF2BPL* was detected on whole exome sequencing (WES), NM_024496.3, c.562C>T (p.R188X), confirmed by Sanger sequencing.

### Subject #3

This is a 20-year-old male who met gross and fine motor milestones early in life. Overall development through the first several years was felt to be normal, although some balance problems with walking may have occurred around 2 years. However, he started stumbling and falling at school at approximately 5-6 years. He developed ataxia and began drooling. A neurological evaluation at age 7 years found nystagmus, hyperactive reflexes, bilateral Babinski signs, and dysmetria. Worse leg weakness and more falls were reported at 8 years. By age 10 his symptoms had progressively worsened, and he had begun experiencing 1-3 myoclonic jerks daily, feeding difficulties, facial weakness, and had lost his speech and ability to walk. By age 13 he was G-tube dependent (from worsening dysphagia) and had slowed saccadic eye movements and severe contractures of the hands and feet. He could not hold objects. He is wheelchair dependent with extremely limited range of motion of all joints. Spastic quadriplegia is managed with a baclofen pump. He has a roving gaze and no visual attention.

Previous evaluations have included MRI of the brain that was normal at age 7 years, but had diffuse cerebral atrophy with *ex vacuo* dilatation of the lateral ventricles but no evidence of white matter disease at 13 years. At 20 years of age, brain MRI demonstrated severe cerebral volume loss of the bilateral hemispheres, patchy periventricular subcortical white matter signal hyperintensity on fluid-attenuated inversion recovery, and marked thinning of the corpus callosum (Figure 2A). Neuropsychological evaluation with the Vineland Adaptive Behavior Scales (3rd edition) at age 20 years indicated continued global and significant delays across all aspects of the patient’s functioning. A *de novo* nonsense variant in *IRF2BPL* was identified on WES, NM_024496.3, c.562C>T (p.R188X) and Sanger confirmed.

**Figure 2:**
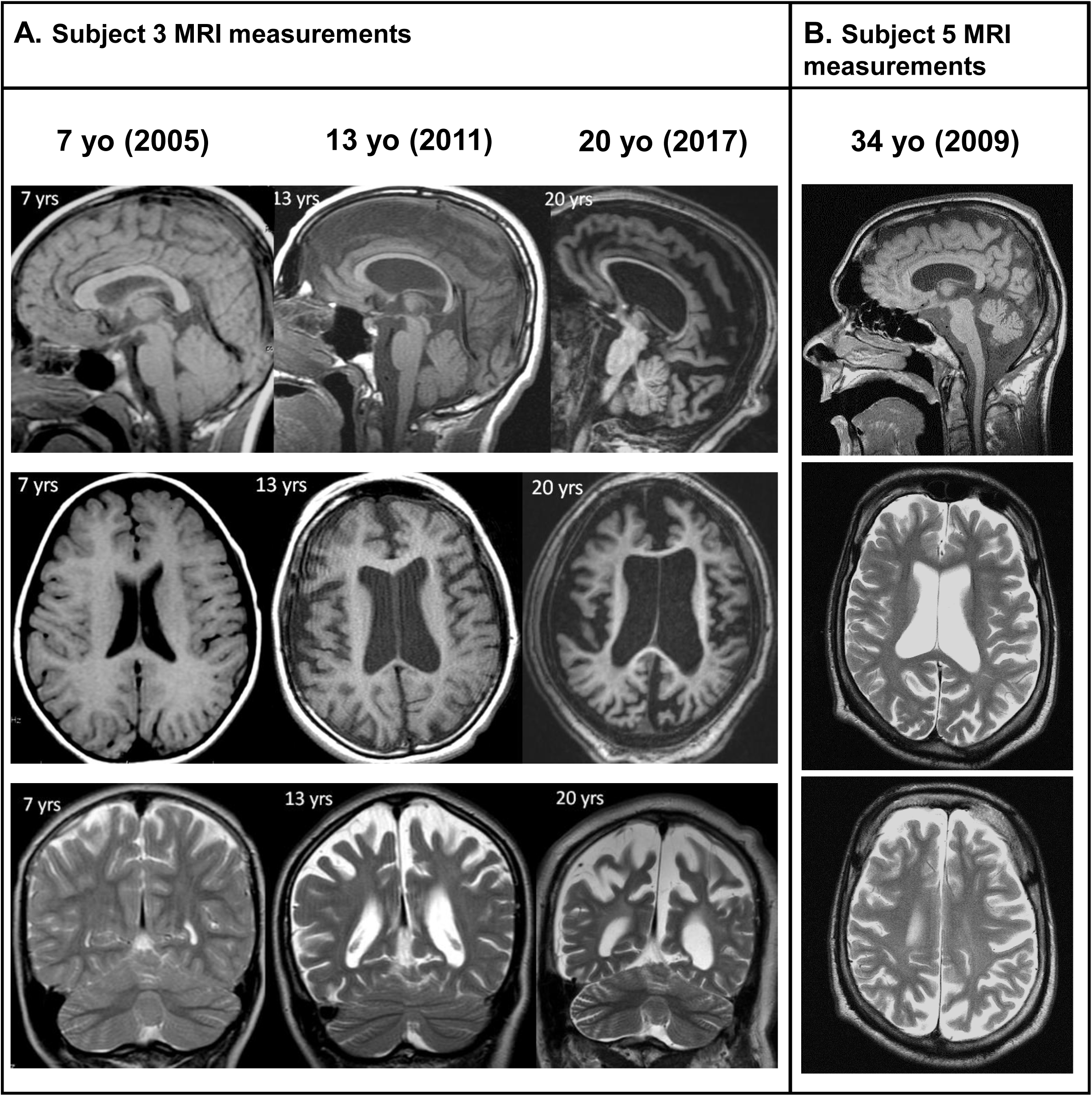
Progressive cerebral atrophy in patients with *IRF2BPL* nonsense truncations. (A) Brain MRI for subject 3 at 7, 13, and 20 years (top row, axial FLAIR; bottom row, sagittal T1). Brain MRI was normal at 7 years. However, at age 13, there was severe diffuse cerebral atrophy with *ex-vacuo* dilatation of the lateral ventricles. There may be slightly increased white matter signal in the peritrigonal region, but otherwise the white matter appears intact and does not suggest a leukodystrophy. The cerebellum has only minimal atrophy. There is mild atrophy of the basal ganglia and brainstem (not shown). At age 20 years, there is further atrophy, including severe volume loss in the bilateral cerebral hemispheres, further thinning of the corpus callosum, mild worsening of the increased white matter signal in the peritrigonal region, further atrophy of the cerebellum and brainstem. (B) MRI images of subject 5 at 34 years depicting global cerebral and cerebellar atrophy, thinning of corpus callosum and brainstem without focal brain lesions (axial T2-weighted and sagittal T1-weighted). Images were taken on a 1.5T Siemens.

### Subject #4

This is a 16-year-old female with global developmental, hypotonia, and speech delay. She had prenatal exposure to alcohol and controlled substances. She was diagnosed with subclinical epilepsy, due to staring episodes and abnormal EEGs, wide-based gait with high-arched feet at 6 years. Cognitive testing was completed at this age (Bracken Basics Concepts III Receptive and School Readiness Composite; Conner’s Parent Rating Scale; Child Behavior Checklist) and resulted in a diagnosis of ADHD and intellectual functioning at approximately the 1-3 year old level. Since 12 years old, there has been a decline in gross and fine motor abilities, speech and self-help skills with a recent diagnosis of catatonia and lower extremity dystonia in the past year. This was initially attributed to sexual abuse, but has been persistent since she was removed from the abusive environment. She had a normal brain MRI at 6 and 13 years of age. At 15, a thin corpus callosum, still within normal limits, was observed on brain MRI. A swallow study at 15 years of age revealed silent aspiration with all tested consistencies. She now is gastrostomy tube dependent, and requires assistance with all activities of daily living. She is largely wheelchair dependent, though able to walk with assistance for short periods. She has progressive macular degeneration that was stable upon her latest examination at 15 years.

Biochemical and cytogenetic testing were completed and are summarized in Supplementary Table 2. She had exome sequencing in 2016 that revealed a mutation in *BEST1* that explains her retinal symptoms (Macular dystrophy, vitelliform, 2, VMD2 [MIM: 153700]). A truncating variant in *IRF2BPL*, NM_024496.3, c.379C>T (p.Q127X) was also reported and was Sanger confirmed.

### Subject #5

This is a 43-year-old male who had psychomotor retardation with hypotonia. He had febrile seizures between 2 and 4 years of age, was diagnosed with epilepsy at 10 years, and developed refractory myoclonic epilepsy by 12 years of age. His neurodevelopmental course showed regression, and at the age of 15 years he had ataxia, spastic rigidity, dystonia and dyskinesia, and a diagnosis of spastic-athetoid cerebral palsy. Progressive cognitive deterioration was noticed, and communication became difficult. At 28 years, he could barely walk independently and could not feed himself. Together with the refractory seizures, the progressive motor problems quickly led him to be wheelchair-bound. At 29 years, he had a vagal nerve stimulator placed to control his seizures, and an intrathecal baclofen pump was placed at 30 years to treat his spasticity. Dystonic attacks involving the axial and appendicular musculature became more frequent. He was treated for recurrent aspiration pneumonia and spontaneous pneumothorax. At the age of 35, his epilepsy was under control, but his dystonia became generalized (limbs, axial, facial with tongue protrusion) and worsened gradually, leading to a completely bedridden state. Treatment with botulin toxin gave only limited benefit. At 42 he had a right-sided pallidotomy, also with limited and transient beneficial effect. Brain MRI at 34 showed global atrophy (cerebral, cerebellar and brainstem), thinning of corpus callosum, and no white matter or cortical lesions (Figure 2B). MR-spectroscopy showed normal findings.

Biochemical, cytogenetic and molecular analyses are summarized in Supplementary Table 2. Trio WES at Ghent University Hospital revealed a *de novo* heterozygous nonsense mutation (NM_024496.3, c.376C>T (p.Q126X)) in *IRF2BPL*, resulting in a premature stop codon, which was Sanger confirmed.

### Subject #6

This is an 11-year-old male with gross motor and speech delays. He rolled over at 14 months and walked at 21 months. At approximately 14 months he received a diagnosis of generalized myoclonic epilepsy that was difficult to control. He had complete regression of speech at 2 years and remains nonverbal, though his receptive language skills appear to be appropriate. He was diagnosed with autism spectrum disorder (ASD) at 3.5 years. He has not had additional regression in developmental skills.

Evaluations are summarized in Supplementary Table 2 and include a normal brain MRI at 7 years of age. Trio WES detected a *de novo* missense variant in the candidate gene *IRF2BPL*, NM_024496.3, c.1115C>G (p.P372R) that was confirmed by Sanger sequencing.

### Subject #7

The patient is a female who was diagnosed with infantile spasms at age 6 months. Seizures subsided with combination antiepileptics at approximately 9 months. Anti-seizure medications were discontinued thereafter with no recurrence of episodes. Brain MRI and MR spectroscopy were both normal. She met gross motor milestones early in life, though on last assessment at 17 months, she continued to be non-verbal and hypotonic. Providers noted deceleration of developmental progress with onset of infantile spasms at 6 months of age, though the patient has not had frank regression. Facial examination revealed subtle dysmorphic facial features including epicanthal folds, mild telecanthus, and almond-shaped eyes with round facies. Developmental assessment using the Bayley Scales of Infant and Toddler Development, 3rd edition (Bayley-3) at 15 months revealed cognitive, expressive and receptive language, and fine and gross motor functioning below levels expected for age (< 1st percentile). There was no family history of seizures or developmental delay. A chromosomal microarray revealed a 400 kb interstitial duplication at 7q31.31 of uncertain clinical significance. Additional tests are listed in Supplementary Table 2. A *de novo* missense variant was detected on *IRF2BPL*, NM_024496.3, c.1254G>C, (p.K418N) on trio WES on 42/87 of the reads.

## Methods

Exome sequencing: Subjects had WES performed on a clinical or research basis. Exome data are summarized in Supplementary Table 3. Across the performing labs, the minimum average depth of coverage was 100X across assays, and minimum proportion of the target at >10X coverage was 95%.

Exome reanalysis (subject 1): FASTQ files were obtained from the relevant source with parental consent. Alignment and variant calling have been previously described.^34^ Novel genotypes were filtered for quality and control observations in public database controls. We highlighted variants in known OMIM genes or mouse essential genes, loss-of-function (LoF) variants that are in genes with known pathogenic LoF variants or reported as haploinsufficient by ClinGen^35^, and LoF intolerant by high probability of LoF intolerance (pLI) score (>0.9).^20^ Conservation of the variant site is reported with the Genomic Evolutionary Rate Profiling (GERP) score.^36^

Generation of fly stocks: All fly strains used in this study were generated in house or obtained from the Bloomington Drosophila Stock Center (BDSC) and cultured at room temperature unless otherwise noted. The *pits*^*MI*02926-*TG4*.1^ allele was generated by genetic conversion of the MiMIC^37,38^ (Minos Mediated Integration Cassette) insertion line, *y*^*1*^ *w*^*\**^ *Mi*{*MIC*}*CG11138*^*MI02926*^ via recombination-mediated cassette exchange (RMCE) as described^39-41^. The recessive lethality associated with the *pits*^*MI02926-TG4.1*^ allele was rescued using an 80 Kb P[acman] duplication (*w[1118]; Dp*(*1;3*)*DC256, PBac*{*y[+mDint2] w[+mC]=DC256*}*VK33*)^42^ as well as by a 20 kb genomic rescue construct (see below). Expression pattern of *pits* was determined by crossing *pits*^*MI02926-TG4.1*^ to *UAS-mCD8-eGFP* (BDSC_32184). In addition, the *y*^*1*^ *w*^*\**^ *Mi*{*MIC*}*CG11138*^*MI02926*^ line was converted to a protein trap line (*pits*::GFP) by injection of a construct that could either produce a splice acceptor (SA)-T2A-GAL4-polyA mutant allele or a SA-eGFP-splice donor (SD) protein trap allele depending on the inserted direction. The successful generation of the *pits*::GFP allele was confirmed by both PCR of genomic DNA and anti-GFP immunohistochemistry/Western blotting of fly tissue.

All transgenic constructs were generated by Gateway (Thermo Fisher) cloning into the pUASg-HA.attB plasmid.^43^ The human *IRF2BPL* cDNA clone^30^ was made to match the NM_024496.3 transcript. Flanking Gateway *attB* sites were added by PCR of the template cDNA clone, and then shuttled to the pDONR223 by BP clonase II (Thermo Fisher). Variants were generated by Q5 site directed mutagenesis (NEB), fully sequenced (Sanger) and finally cloned into pUASg-HA.attB via LR clonase II (Thermo Fisher). All expression constructs were inserted into the VK37 (*PBac*{*y[+]-attP*}*VK00037*) docking site by ϕC31 mediated transgenesis.^44^ The 20Kb genomic rescue (GR) line was generated by inserting the P[acman] clone CH322-141N09 (BACPAC Resources)^45^ into the VK37 docking site. The *UAS-pits-RC* flies were generated by obtaining the *pits* cDNA (RE41430, RC isoform) clone from the Drosophila Genomics Resource Center (DGRC) and performing Gateway (Thermo Fisher) cloning into pUASg-HA.attB. The pits RNAi line (*P*{*KK108903*}*VIE-260B*) was obtained from Vienna Drosophila Resource Center (VDRC). The GAL4 lines used in this study from BDSC are: *nSyb-GAL4* (*y*^*1*^ *w*^*\**^; *P*{*w*^*+m\**^*=nSyb-GAL4.S*}*3*) BDSC_51635, *Rh1-GAL4* (*P*{*ry*^*+t^7^.^2^*^*=rh1-GAL4*}*3, ry*^*506*^) BDSC_8691, *Act-GAL4/CyO* (*y*^*1*^ *w*^*\**^; *P*{*w*^*+mC*^*=Act5C-GAL4*}*25FO1/CyO, y*^*+*^) BDSC_ 4414.

*Drosophila* behavioral assays: To perform the bang sensitivity assay^46^, flies were anesthetized no sooner than 24 hours prior to testing with CO_2_ and housed individually. At the time of testing, flies were transferred to a clean vial without food and vortexed at maximum speed for 15 seconds. The time required for flies to recover in an upright position was measured. Typically 25 flies were tested per data point. The climbing assay^47^ was performed in a similar manner but at the time of testing flies were transferred to a fresh vial and given 1 minute to habituate before tapped to the bottom of the vial three times and examined for negative geotaxis (climbing) response to reach the 7 cm mark on the vial. Flies were given a maximum of 30 seconds to reach the top.

Histological and ultrastructural analysis of the fly retina: 45-day-old flies were dissected in ice cold fixative (2% PFA / 2.5% glutaraldehyde / 0.1 M sodium cacodylate). Heads were fixed, dehydrated and embedded in Embed-812 resin (Electron Microscopy Sciences) as described previously.^48^ For histological sections (200 nm thick) were stained with toluidine blue and imaged with a Zeiss microscope (Axio Imager-Z2) equipped with an AxioCam MRm digital camera. For transmission electron microscopy (TEM), sections (50 nm thick) were stained with 1% uranyl acetate and 2.5% lead citrate. TEM images were obtained using a transmission electron microscope (model 1010, JEOL). Images were processed with Photoshop (Adobe) and 45 day old toluidine blue images were color matched to the 5 day old images for clarity.

Overexpression of human *IRF2BPL* in *Drosophila*: We overexpressed reference and variant *IRF2BPL* cDNAs in flies by crossing the *UAS-IRF2BPL* males to virgin female flies from a ubiquitous (*Act-GAL4*) driver stock. The progeny of these crosses were cultured at various temperatures (18°C, 22°C, 25°C, and 29°C) to express the human proteins at different levels (lowest expression at 18°C and highest expression at 29°C).^38,49^ Expected Mendelian ratios of flies was assessed. For temperatures ≥22°C over 100 flies were assessed for each cross. For 18°C crosses, over 50 flies were assessed from each group.

For additional methods please see supplemental material.

## Results

### Summary of Clinical Findings

The subjects described here are between 2.5 and 43 years of age, including one who died at 15 years of age. Comprehensive clinical information is included in Supplementary Table 1 and Supplementary Table 2.

All seven individuals have monoallelic variants in *IRF2BPL* (Table 1 and Supplementary Table 1). The nonsense variants identified are: p.Q126X, p.Q127X, p.E172X, and p.R188X (found in 2 individuals). There was remarkable similarity in the clinical course of the five cases that had nonsense variants. These include an unremarkable course in their initial development, followed by loss of developmental milestones and development of a seizure disorder at a variable age, with 2 years being the youngest. Additional findings included a movement disorder with dystonia and choreoathetosis, and cerebellar signs such as ataxia, dysarthria, dysmetria and dysdiadochokinesia. Three of the probands had had anomalies of eye movements but had normal retinae and optic nerves. Head circumference appeared to remain appropriate for age in those with available measurements. Brain MRI was normal early in life, but cerebral and cerebellar atrophy was noted in the two oldest cases. Cognitive status was difficult to ascertain, particularly as verbal skills were lost. However, cognitive abilities appeared to remain intact over a short follow up period in one case and to deteriorate in an older individual. One case was deceased at 15 years of age when the family elected to limit care after continued decline. The variants were *de novo* in all for whom parental testing was available (three out of five cases).

Missense variants (p.P372R and p.K418N) were also reported in *IRF2BPL* for two individuals who had a variable phenotype of global developmental delay and seizures. The older case (subject 6) had a diagnosis of ASD. Despite the frequent finding of seizures, antiepileptic drugs were successfully weaned in subject 7, without recurrence over a short follow up period. In both missense cases, the variants were *de novo*. Neither case has developed a progressive loss of milestones, abnormal eye movements, or a movement disorder at 2 or 10 years of age.

### *IRF2BPL de novo* variants are deleterious based on bioinformatics data

*IRF2BPL* is a gene that is highly intolerant to variation with an Residual Variation

Intolerance Score (RVIS)^50^ of 9.3% and a pLI score of 0.97, with no observed LoF variants included in the calculation.^20^ The only LoF variants present in ExAC and gnomAD are frameshifts, which either do not pass quality filtering or appear to be artifactual calls in repetitive regions using the browser-visualization tool.^20^ *IRF2BPL* is also constrained to missense variants (z = 4.73).^20^ All variants found in the subjects were absent from the gnomAD and ExAC databases. The Combined Annotation Dependent Depletion (CADD)^51^ score for all of the variants was > 34, which places them among the variants predicted to be most deleterious. The four nonsense variants introduce a premature termination codon either downstream (p.E172, p.R188) or at the end (p.Q126, p.Q127) of the poly-glutamine tract and upstream of the first PEST sequence (Figure 1A), whereas the missense variants are within the variable region of the protein. We used the DOMINO tool to assess the likelihood of monoallelic variants in *IRF2BPL* to cause a Mendelian disease. IRF2BPL had a DOMINO score of 0.962 out of 1 predicting that monoallelic variants would very likely cause disease (Figure 1B).^52^

### *IRF2BPL* variants are severe loss-of-function mutations based on functional assays in flies

To validate the functional consequences of these variants, we utilized *Drosophila melanogaster* as a model organism. Experiments in fruit flies have previously provided experimental support in identifying causal variants for human disease^19,53-55^ and these approaches have been an integral part of the UDN. ^13,14,16,21,56^ The fly ortholog of *IRF2BPL* is a poorly characterized gene, *CG11138*. This gene was studied in the context of epigenetic regulation through biochemical methodologies during embryogenesis, and was named *pits* (*protein interacting with Ttk69 and Sin3A*).^57^ Although the overall identity (30%) and similarity (36%) between IRF2BPL and Pits may not seem high, the architecture of the protein is very similar and the sequences of the annotated domains show high conservation (79% identity for the zinc finger domain, 76% identity for the C3HC4 RING domain) (Figure 3A). The Pits protein also has a DIOPT^58^ score of 12/15 suggesting that it is likely to be a true ortholog of IRF2BPL. Two other human paralogs of IRF2BPL share a similar high DIOPT score (12/15 for *IRF2BP1*; 11/15 for *IRF2BP2*). These data indicate that *pits* is the sole fly gene that is orthologous to the three IRF2BP family genes in humans.

**Figure 3:**
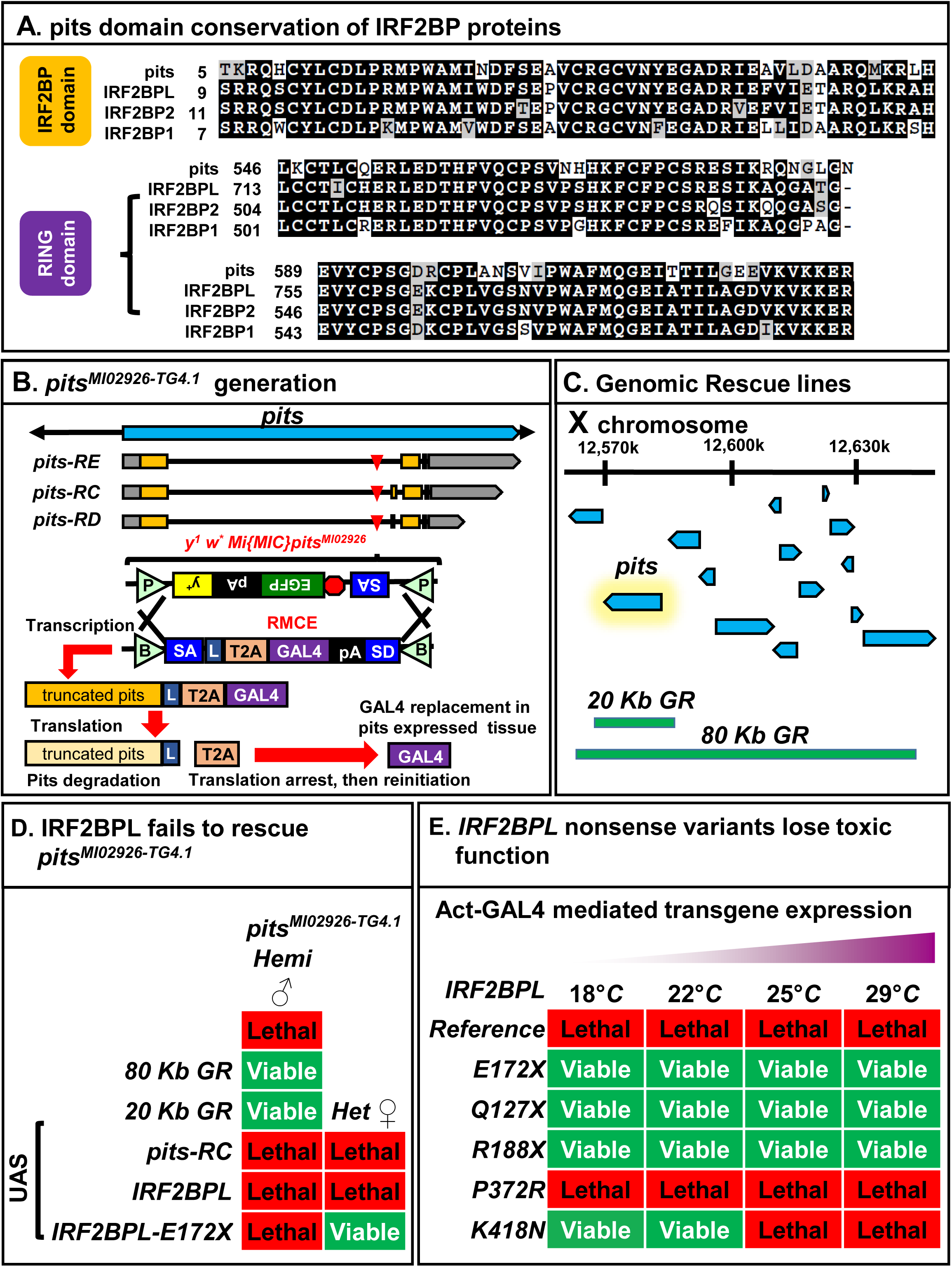
The IRF2BPL homolog, Pits, is highly conserved, and human disease variants display dramatic loss-of-function. (A) The two annotated domains (IRF2BP zinc finger and C3HC4 ring) for IRF2BPL (as well as IRF2BP1 and IRF2BP2) display very high conservation with the fly homolog Pits. (B). The *pits*^*MI02926-TG4.1*^ allele was generated by genetic conversion of *y*^*1*^ *w*^*\**^ *Mi*{*MIC*}*CG11138*^*MI02926*^ by recombination-mediated cassette exchange (RMCE) *in vivo*. The resulting mutant incorporates a SA-T2A-GAL4 that acts as an artificial exon resulting in early truncation of the *pits* transcript and cellular replacement with expression of GAL4 under the endogenous pits regulatory elements. (C) Genomic location of genomic rescue (GR) constructs inserted on chromosome 2 (VK37) of the fly. Note that the 20 Kb rescue line is specific to only the *pits* gene. (D) Reintroduction of either GR construct (Figure 3C) rescues lethality for *pits*^*MI02926-TG4.1*^ flies but rescue is not observed by overexpression of the fly or human cDNA. Female *pits*^*MI02926-TG4.1*^/*FM7* virgins are crossed to males of either GR or UAS lines, and progeny are examined for males containing *pits*^*MI02926-TG4.1*^ and the rescue construct (minimum progeny examined n=91). Examination of female flies heterozygous for the presence of *pits*^*MI02926-TG4.1*^ and the rescue construct reveals a lack of toxicity in female *pits*^*MI02926-TG4.1*^ */+; UAS-IRF2BPL-E172X/+* flies indicating loss-of-function. (E) The ubiquitous expression of *UAS-IRF2BPL* or variants with *Act-GAL4* reveals that all nonsense variants are strong loss-of-function mutations and the p.K418N causes partial loss of function.

To generate a *pits* mutant fly, we used a MiMIC insertion in an intron of *pits*, named *Mi*{*MIC*}*CG11138*^*MI02926*^ (Figure 3B).^37,38^ MiMICs are engineered transposable elements that contain inverted *attP* sites derived from phage ΦC31 flanking a swappable cassette. This allows replacement of the content of the *MiMIC* insertion using Recombination-Mediated Cassette Exchange (RMCE) by expressing the ΦC31 integrase to swap the *MiMIC* cassette with a *SA-T2A-GAL4-polyA* (T2A-GAL4) cassette.^39-41^ This insertion results in a truncated *pits* transcript due to the polyA signal. During translation this short transcript produces a short protein that is truncated at the T2A site yet allows re-initiation of translation to produce the GAL4 protein. The GAL4 protein is expressed in a proper spatial and temporal fashion, i.e. those of the endogenous gene^38^ (Figure 3B). This allows rescue of the *T2A-GAL4* induced allele, typically a null allele, with a UAS-(fly)cDNA for about 70% of the genes tested.^40^

The *pits* gene is on the X chromosome, and *pits*^*MI02926-TG4.1*^ males are hemizygous lethal, as they fail to survive past the first instar larval stage, and most die as embryos. Lethality can be rescued to viable adults by introduction of an 80 Kb or a 20 Kb P[acman] genomic BAC rescue (GR) construct, the latter only carrying the *pits* gene^42,44^ (Figure 3C and 3D). Hence, *pits* is an essential gene, and expression of the gene in the proper genomic context fully rescues the LoF of *pits.*

We attempted to rescue lethality observed in *pits*^*MI02926-TG4.1*^ flies by overexpression of UAS-*pits* or UAS-*IRF2BPL* and failed to obtain viable flies. Intriguingly, we were also unable to obtain viable heterozygous female flies that contain both the *pits*^*MI02926-TG4.1*^ allele and UAS-*pits* or *UAS-IRF2BP* (Figure 3D). These data show that overexpression of the fly or human IRF2BPL in the cells that endogenously express Pits is toxic to the fly. Indeed, expression of the *UAS-IRF2BPL* under the control of a ubiquitous driver (*Act-GAL4*) also causes lethality (Figure 3E).

The above data show that the lethality caused by overexpression of the reference *IRF2BPL* gene can be used as a functional assay to test if the variants are functional (toxic) as well. To modulate the levels of expression we ubiquitously expressed the variants with *Act-GAL4* as the UAS-cDNA expressed using this driver exhibits temperature dependence, with significantly greater expression at higher temperatures.^38,49^ We conducted these experiments at 18°C, 22°C, 25°C and 29°C. Expression of the reference *IRF2BPL* cDNA consistently causes lethality at all temperatures tested, whereas the three nonsense variants (p.E172X, p.Q127X, and p.R188X) consistently produced viable animals at all temperatures tested (Figure 3E). This provides evidence that the truncated proteins are not toxic and are very likely LoF alleles. Interestingly, the missense variant p.K418N is lethal when expressed at higher temperatures, whereas the flies are viable at lower temperatures, indicating that it retains some toxic function and hence behaves as a partial LoF allele. The p.P372R variant still remained lethal upon ubiquitous expression, even at lower temperatures indicating its toxic function is similar to the reference protein at least in this assay. We confirmed that the UAS-driven human cDNA in flies expresses IRF2BPL at relatively similar levels by making HA-tagged constructs for the reference and p.E172X truncation and performing Western blot using an anti-HA antibody, as commercially available antibodies for IRF2BPL recognize epitopes downstream of the premature termination codon, in the C-terminus of IRF2BPL. UAS expression of these constructs in neurons by *nSyb* (neuronal Synaptobrevin)*-GAL4* was relatively similar (Supplemental Figure 2A), Additionally, we confirmed that the untagged reference and two missense variants were expressed at similar levels using a commercial antibody against IRF2BPL and performing Western blot, suggesting that the reduced toxicity of p.K418N is not due to destabilization of the protein (Supplemental Figure 2B).

### *pits* is expressed and required in the CNS

To examine the endogenous levels of *pits* as well as its subcellular localization, we generated a GFP protein trap line of *pits*. We integrated a protein trap (splice acceptor (SA)-linker-eGFP-linker-splice donor (SD)) cassette into the *pits*^*MI02926*^ via RMCE. Although the SA-eGFP-SD functions as an artificial exon in the middle of the *pits* gene (Supplemental Figure 3A), *pits::GFP* flies are homozygous viable and do not display any obvious phenotype indicating that the internal GFP tag is not deleterious. This is in concert with our previous findings that most (75%) proteins tolerate the presence of an internal GFP incorporated by the SA-eGFP-SD MiMIC insertions.^38^ The GFP fusion proteins also reflect the localization of the endogenous proteins.^37,38^ We first confirmed the presence of a single protein of the appropriate size on SDS-PAGE in males (Figure 4A). As shown in Figure 4B and 4C, the tagged protein is widely expressed in the brain. In the third instar larval brain the protein is widely expressed and is enriched in the mushroom body (MB, Figure 4B, yellow arrow). In the adult brain Pits::GFP is localized in most neurons, including the cell bodies and nuclei (co-staining with a pan-neuronal nuclear marker, Elav) of many neurons as well as their axons (Figure 4C). This is in agreement with a previous study showing *pits* is expressed in the nucleus.^59^ Pits::GFP does not show expression in the glia of the adult CNS upon pan-glial staining with Repo (Supplemental Figure 3B). Additionally, we also determined the expression of pits using the GAL4 reporter from *pits*^*MI02926-TG4.1*^ allele crossed to a UAS-membrane bound GFP (Supplemental Figure 3C) which allow detection of cells that express the gene of interest at low levels ^40^ We also generated z-stacks confocal image movies of pits::GFP co-stained with Elav (Supplemental Movie 2) and Repo (Supplemental Movie 3). These data reveal that *pits* is highly expressed in the MB, the learning and memory center of the flies^60^ as well as in the antennal mechanosensory and motor center, which are required for balance and hearing and motor coordination, a cerebellar insect equivalent.^61^ Again, no expression was observed in glia from these analyses.

**Figure 4:**
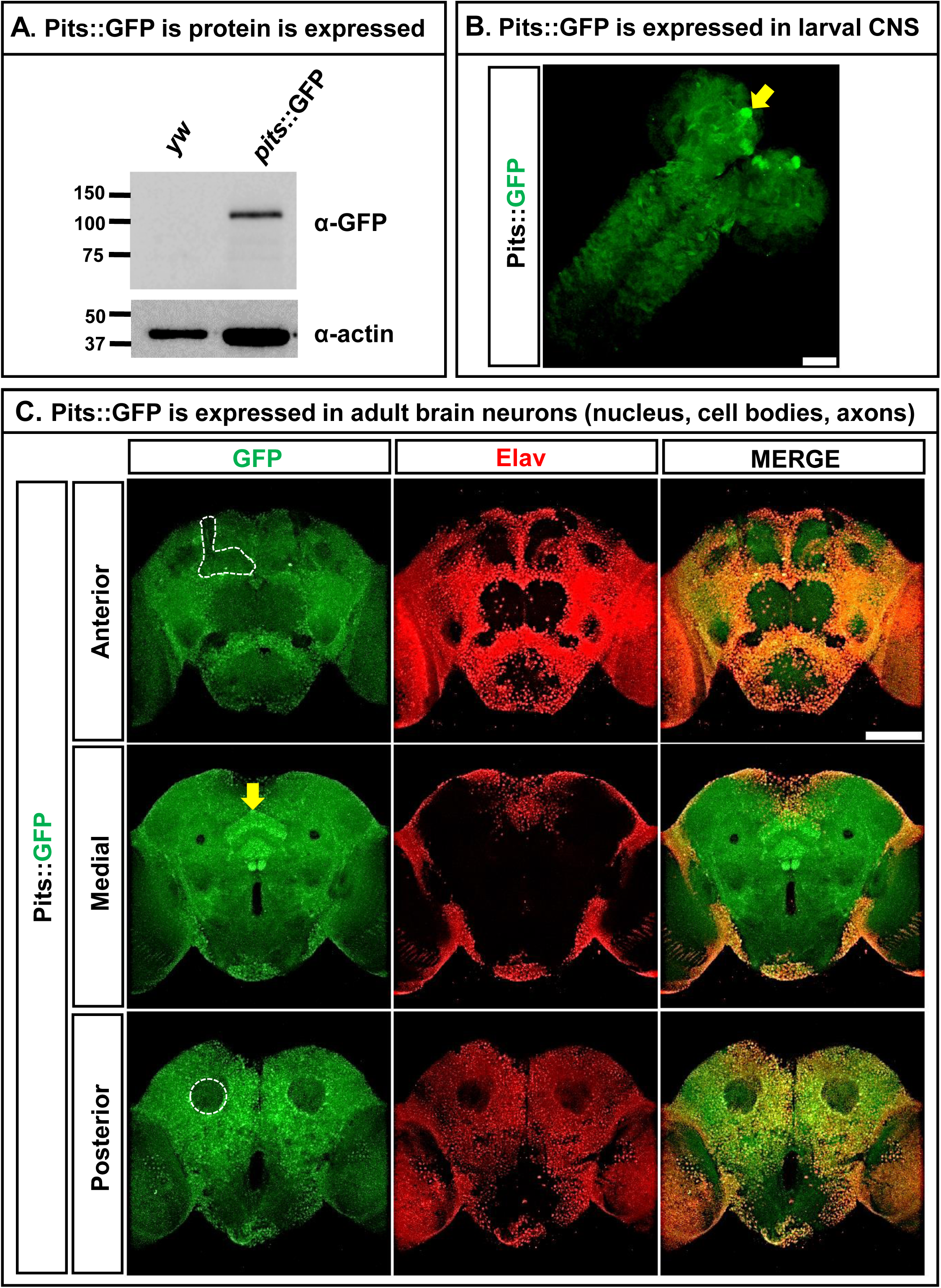
The *IRF2BPL* homolog, *pits*, is expressed in both the developing and adult CNS and is present in the nucleus of a wide subset of neurons. (A) Male fly heads were lysed and run on SDS-PAGE to determine the presence of Pits::GFP. (B) Pits is widely expressed in 3rd instar larvae, assessed by immunostaining of homozygous pits::GFP animals and viewed by confocal microscopy (z-stack - max projection). Note the enrichment in the mushroom body (yellow arrow). (C) Single slice confocal images of the adult CNS show *pits* expression in neurons (co-localized with Elav). Notably *pits* is expressed in the cell bodies of the adult mushroom body (top left panel), is enriched in the central complex (yellow arrow) and is not present in the dendrites of the mushroom body (bottom left panel). Scale bar is 50 μm.

Due to the lethality observed in *pits*^*MI02926-TG4.1*^ flies, we determined the function of pits in the brain through gene knock-down studies using RNA interference (RNAi).^62^ Ubiquitous knock-down of pits using *Act-GAL4* resulted in semi-lethality (Figure 5A). This RNAi has specificity to pits as it consistently reduces the endogenous Pits protein level to ∼50% of control RNAi when assessed via Western blot (Figure 5B). To determine if partial knock-down of pits results in neurological defects, we expressed the pits RNAi with the *nSyb-GAL4* driver. Young flies (∼5 days post-eclosion) displayed normal climbing (Figure 5C) and mechanical stress tolerance (bang sensitivity) (Figure 5D). However, aged flies that were 30 and 45 days post-eclosion displayed progressive abnormalities in climbing, and became bang sensitive at 30 days post-eclosion when compared to controls (Figure 5C and 5D).

**Figure 5:**
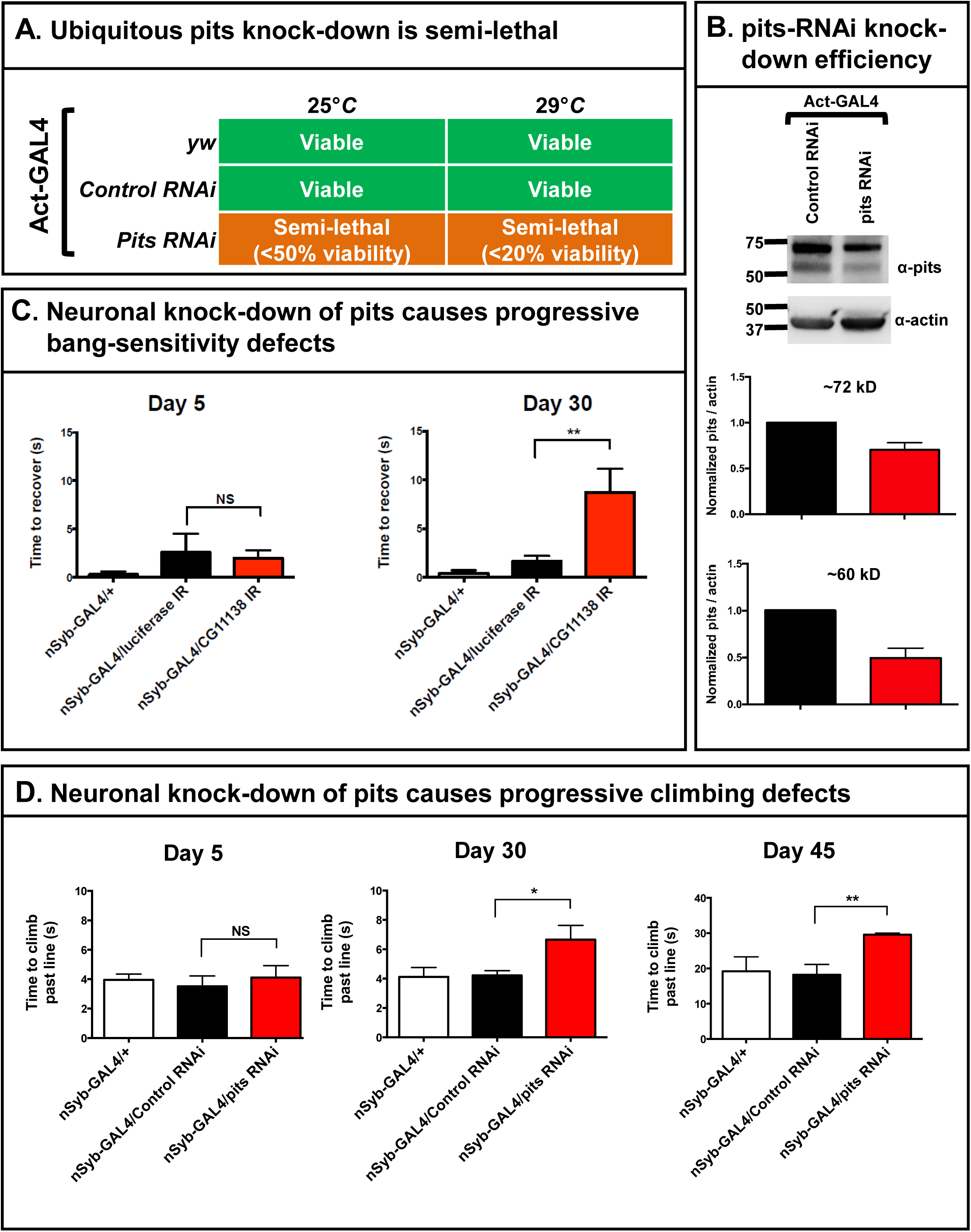
Neuronal knock-down of *pits* leads to progressive behavioral deficits. (A) Ubiquitous knock-down of *pits* results in semi-lethality, as shown by lower than expected genotypic ratios of survival into adulthood. Flies were compared to *Act-GAL4>control-RNAi* and *y w Act-GAL4/+*. (B) The *pits* RNAi can partially knock-down ∼50-60% of the *pits* isoforms consistently observed in female flies. (C) Pan-neuronal knock-down of *pits* (*nSyb-GAL4>pits-RNAi*) leads to a bang-sensitive paralytic phenotype in aged animals that is not observed in control flies or young animals. Multiple cohorts of flies were anesthetized and single housed 24 hours prior to testing with 15 seconds of vortexing in an empty vial. Statistical analyses were with one-way ANOVA followed by Tukeys post-hoc test. Results are means ±SEM (*p<0.05; NS, not significant) (D) Pan-neuronal knock-down of *pits* using RNAi leads to progressive climbing deficits that are only observed in aged flies. Singly housed flies (similar to above) were given 1 minute to habituate to an empty vial before being tapped 3 times. Flies were given 30 seconds to cross the 7 cm mark on the vial. Flies that failed to cross the line were given a score of 30 (only seen at day 45). One-way ANOVA followed by Tukeys post-hoc test. Data are mean ±SEM (*p<0.05, **p<0.01; NS, not significant).

To determine if an age-dependent deterioration in neural morphology can be observed, we reduced *pits* expression in photoreceptors of the fly eye using rhodopsin (*Rh1*)-*GAL4*. The fly retina is a well-characterized system to explore neurodegeneration in *Drosophila* due to the highly stereotypical organization of neurons (photoreceptors) and glia cells (pigment cells, cone cells) in this tissue^19,63^. Indeed, Pits::GFP shows a robust nuclear signal in the photoreceptors of the fly retina when co-stained with Elav (Supplemental Figure 4). Flies were raised in a 12 hr light/dark cycle for 45 days, and histological examination of retina sections stained with toluidine blue was performed. We observed a severe disorganization of the ommatidia (units of photoreceptors) and notable rhabdomere (light-sensing organelle) loss (Figure 6A). This loss was not observed in young flies (5 days post-eclosion), nor in aged control RNAi flies. To examine ultrastructure, we performed transmission electron microscopy (TEM) of the same tissue and observed numerous abnormalities in the ultrastructure of the *Rh1-GAL4>pits-RNAi* retina compared to age matched *Rh1-GAL4>control-RNAi* eyes (Figure 6B and 6C, Supplemental Figure 5 and Supplemental Figure 6): 1) a significant decrease in intact rhabdomeres (Figure 6D); 2) a significant increase in the presence of tubulovesicular like structures (TVS), associated with some neurodegenerative models^64^ (Figure 6C, red arrow); and 3) the abnormal presence of neuronal lipid droplets in photoreceptors (Figure 6C, yellow arrow, Supplemental Figure 7A), often associated with mutants that cause high reactive oxygen species (ROS) accumulation.^65^ However, we did not detect obvious changes in mitochondrial morphology or the number of mitochondria per photoreceptor (Supplemental Figure 7B). In summary, *pits* is required for proper maintenance of neuronal function and structure in flies.

**Figure 6:**
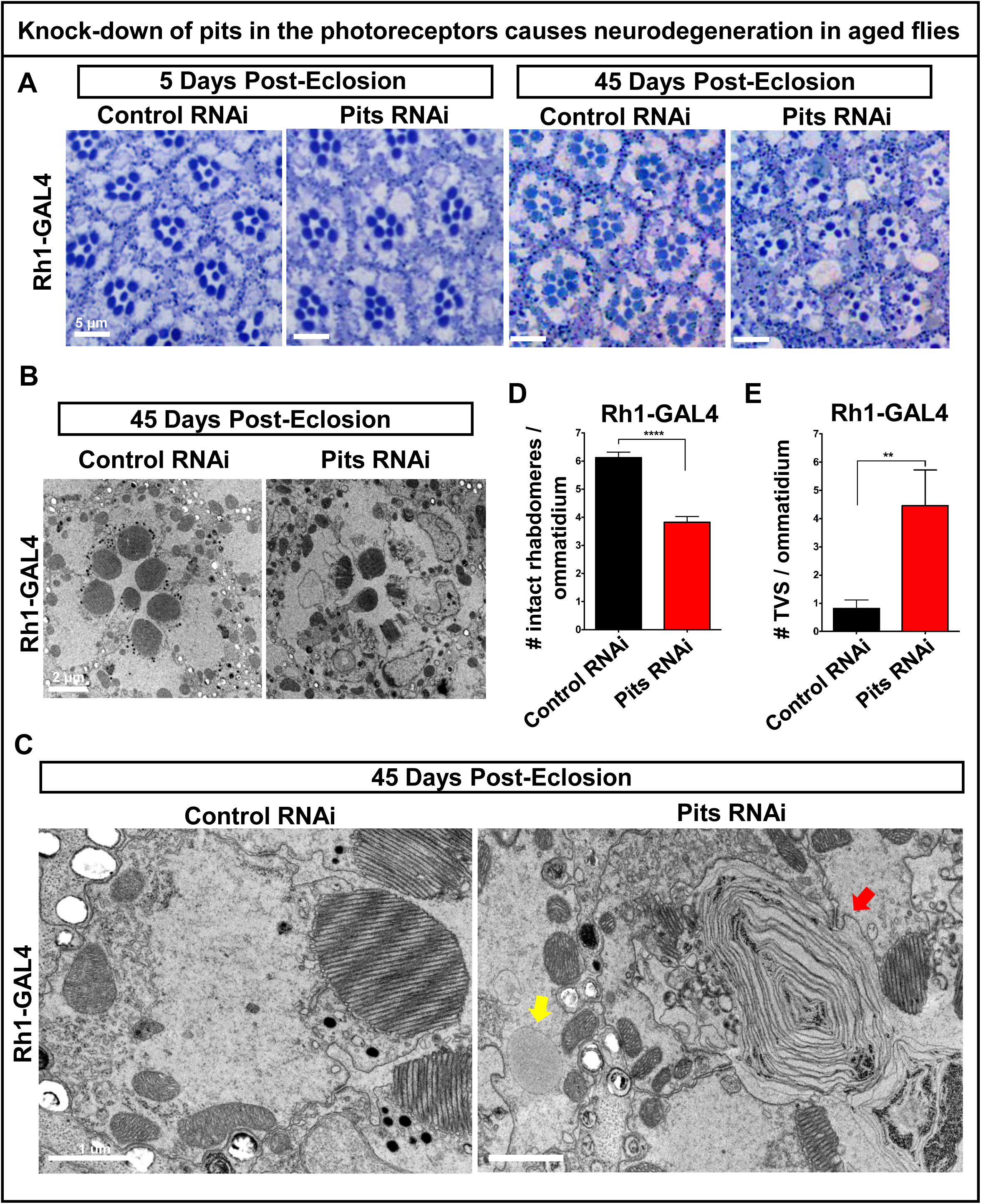
Knock-down of pits in the photoreceptor leads to degenerative phenotypes in aged animals. (A) Toluidine blue staining of 5- and 45-day-old retina from *Rh1-GAL4>luciferase-RNAi* and *Rh1-GAL4>pits-RNAi flies* revealing disorganization of the ommatidtia and photoreceptor loss. Scale bar is 5 μm. (B to C) TEM images showing the photoreceptors and the ommatidia (B, scale bar is 2 μm) and photoreceptors (C, scale bar is 1 μm) section. The red arrow indicates the presence of tubulovesicular like structures (TVS), and the yellow arrow indicates neuronal lipid droplets (C). Further TEM images are available in Supplemental Figures 5 and 6. Quantification of rhabdomere loss (D) and TVS structures (E) on a minimum of n=27 randomized sections from the retina from three animals per genotype. Statistical analyses were with unpaired two-tailed t-tests. Results are mean ±SEM (**p<0.01, ****p<0.0001).

## Discussion

We present seven subjects with rare heterozygous variants in *IRF2BPL*, a gene that has not previously been associated with disease in humans. The cases with nonsense variants exhibited a remarkably similar, progressive course of neurological regression that eventually led to severe disability. In contrast, the missense variants observed in two subjects are associated with milder neurological symptoms such as seizures, developmental delay, and ASD. Interestingly, the nonsense variants in this cohort cluster in or just downstream of the polyglutamine tract within the variable region, while the missense variants map further downstream (Figure 1A). The bioinformatics signature of the gene suggests that *IRF2BPL* is highly intolerant to variation, and that the variants reported in the cohort are among the most deleterious. The DOMINO score was suggestive of dominant inheritance. RNA sequencing from a blood sample for subject 1 (c.514G>T (p.E172X) revealed expression of the transcript (data not shown). The lack of nonsense-mediated decay of *IRF2BPL* transcript with a nonsense mutation is consistent with the fact that this is a single exon gene.^66^ Western blotting on subject samples with nonsense variants could not be pursued due to lack of a commercially-available antibody that recognizes an epitope upstream of the premature truncation.

To date, a role for *IRF2BPL* in humans is limited to an association with developmental phenotypes. For example, *IRF2BPL* has been identified in the top 1,000 genes that are significantly lacking in functional coding variation in non-ASD samples and are enriched for *de novo* LoF mutations identified in ASD cases.^67^ Other large scale sequencing studies have identified *de novo* variants in *IRF2BPL* in ASD (2 individuals) and major developmental disorders (2 individuals).^1,3^ Intriguingly the ASD variants include a missense in the conserved DNA binding domain (p.F30L) and a frameshift near the end of the protein (p.A701fs*65). The developmental disorder cohort also has a missense (p.R391C) and frameshift that similarly terminates near the end of the protein (p.L713Pfs*54).^1,3^ The missense variants in these cases compared to our two cases indicate that the full spectrum of IRF2BPL related phenotypes is still developing. The p.A701fs*65 frameshift, in particular, suggests that truncations near the end of the protein may cause variable phenotypes. Another possibility is the presence of mosacism in these large cohort studies. Our findings offer additional evidence that variants in *IRF2BPL* are implicated in neurological symptoms, and additionally extend the phenotype into neurological regression. Furthermore, our model organism experiments using fruit flies supports an important role for *IRF2BPL* in embryologic development as well as neuronal maintenance. *IRF2BPL* is well-conserved and the fly gene, *pits,* is expressed widely in the nervous system during development and in adulthood. The *pits*^*MI02926-TG4.1*^ LoF mutant fails to survive past early larval stages and mostly die as embryos. Although we were not able to rescue the *pits*^*MI02926-*^ *TG4.1* allele with either the human or fly cDNA, the *pits*^*MI02926-TG4*^ is rescued with a genomic rescue construct specific to *pits,* showing that this chromosome does not carry other lethal or second site mutations. Interestingly, ubiquitous overexpression of *IRF2BPL* or *pits* in flies is toxic, suggesting that the gene is highly dosage sensitive. Many genes implicated in neurodegenerative disorders cause overexpression phenotypes in *Drosophila*.^68,69^ In our experiments, however, overexpression of the nonsense variants was not toxic. These results suggest that the mechanism of disease may be through loss of protein function and haploinsufficiency of *IRF2BPL*. Overexpression of the missense variants showed a range of effects. While overexpression of p.K418N was only lethal at higher temperatures (i.e., higher levels of expression), p.P372R was lethal at any temperature. These results suggest that some of the proteins expressed with the missense variants retained a function that was toxic when overexpressed, or perhaps suggest a gain of function mechanism. Understanding the precise molecular function of Pits/IRF2BPL will allow one to design experiments to test this possibility. In summary, either excess or loss of *pits* or *IRF2BPL* is highly detrimental to survival. Given that expression at low levels (*Act-GAL4* at 18°C) is still toxic suggests that the level of expression of the gene product is highly regulated *in vivo* through mechanisms such as ubiquitination through the PEST domain.

There are compelling parallels in phenotypes between a partial Pits reduction in flies and symptoms observed in patients, which may suggest evolutionary conservation in neuronal mechanisms. 1) Almost all patients had seizures or EEG abnormalities, and a reduction of Pits in neurons leads to a bang-sensitive phenotype in flies. Bang-sensitivity is associated with seizure-like paralysis that has phenotypic as well as genetic parallels with human epilepsy.^70,71^

2) The five patients carrying nonsense variants in *IRF2BPL* displayed progressive motor dysfunction that manifested after early childhood. Similarly, neuronal reduction of Pits in flies caused a progressive decline in climbing ability that was not observed in young flies. 3) The two oldest patients, who had nonsense variants, had evidence of cerebral atrophy in adulthood. Correspondingly, we found that a reduction in pits expression in photoreceptors leads to a slow and age-dependent loss of neuronal integrity. 4) The cerebellar symptoms and cerebellar atrophy may correspond to a requirement for Pits expression in the antennal mechanosensory and motor center. These neurons are required for balance, auditory and motor coordination, a cerebellar insect equivalent ^61^ and Pits is abundantly expressed in these cells. These results support the notion that IRF2BPL / Pits have fundamental roles in central nervous system development and maintenance.

IRF2BPL has similarity to its mammalian paralogs IRF2BP1 and IRF2BP2 and has been shown to interact with IRF2 (Interferon Regulatory Factor 2).^72,73^ However, flies lack an obvious homolog to IRF2, indicating that *pits* and *IRF2BPL* may have additional conserved functions that are currently unknown. Recently, a study in *Drosophila* indicated that the fly homologue, Pits, regulates transcription during early embryogenesis by interacting with a histone deacetylase Sin3A (SIN3A and SIN3B in human) and a co-repressor Tramtrack 69 (no identified human ortholog).^57^ Consistent with previous studies in flies and rats, our data suggest both Pits and IRF2BPL are found predominantly in the nucleus.^59,74^ We showed robust expression in the nucleus of photoreceptors and numerous neurons, but also observed Pits::GFP in axons and cell bodies, and little to no staining in the dendrites of the mushroom body. This could indicate that Pits may have a non-nuclear role in specific subcellular compartments in neurons. Finally, through neuron-specific knock-down experiments in flies, we demonstrate that *pits* is important for function and/or neuronal maintenance over time.

We observed different phenotypes within the cohort based on the type of variant. All four nonsense variants truncate the IRF2BPL protein upstream of the putative nuclear localization signal, the conserved C’ terminal RING domain, and multiple putative PEST sequences. Although little is known about IRF2BPL/Pits, most studies implicate its expression and function in the nucleus.^30,57,59,74,75^. However, the RING domain of IRF2BPL has recently been shown to act as an E3-ligase ubiquitinating β-catenin and supressing Wnt signalling in gastric cancer.^32^ It is currently unclear whether this pathway could be altered in IRF2BPL-associated disease. Finally, the predicted PEST sequences suggest that IRF2BPL is highly regulated and support our data that overexpression of the protein can be detrimental. Toxicity was observed by overexpression of fly or human cDNA constructs in all cells (*Act-GAL4*) or within the cell-types in which pits is endogenously expressed (*pits*^*MI02926-TG4.1*^). While *IRF2BPL* ubiquitous expression or overexpression within the cell types that pits is endogenously expressed caused lethality, we did not observe lethality or any remarkable phenotypes by overexpression of IRF2BPL or the variants when expressed specifically in neurons (*nSyb-GAL4*) (data not shown). Therefore, increased Pits/IRF2BPL protein expression may be detrimental to only certain cells. Our functional assays of nonsense and missense variants support haploinsufficiency and tight control of protein expression and/or turn-over. Future studies will determine IRF2BPL targets for ubiquitination and binding partners that mediate neurological phenotypes.

In summary, we have implicated dominant *de novo* variation in *IRF2BPL* to a new neurological disorder in humans. We observed that patients with nonsense variants in this single exon gene suffer from a progressive and devastating neurological regression, and individuals with rare missense variants also show a spectrum of neurological phenotypes. The bioinformatics signature supports a deleterious effect in IRF2BPL at the gene and variant level. We provide functional analysis in flies to support a loss-of-function model of *IRF2BPL*-associated disease. Future studies examining the mechanism of early death in *pits* mutant flies, as well as molecular mechanisms of neurodegeneration in *pits* knockdown animals, will likely shed light on conserved pathways essential for neurological development and provide guidance to design an effective therapeutic or preventive strategy.

## Consortia

The Undiagnosed Diseases Network co-investigators are David R. Adams, Mercedes E. Alejandro, Patrick Allard, Mahshid S. Azamian, Carlos A. Bacino, Ashok Balasubramanyam, Hayk Barseghyan, Gabriel F. Batzli, Alan H. Beggs, Babak Behnam, Anna Bican, David P. Bick, Camille L. Birch, Devon Bonner, Braden E. Boone, Bret L. Bostwick, Lauren C. Briere, Donna M. Brown, Matthew Brush, Elizabeth A. Burke, Lindsay C. Burrage, Shan Chen, Gary D. Clark, Terra R. Coakley, Joy D. Cogan, Cynthia M. Cooper, Heidi Cope, William J. Craigen, Precilla D’Souza, Mariska Davids, Jyoti G. Dayal, Esteban C. Dell’Angelica, Shweta U. Dhar, Ani Dillon, Katrina M. Dipple, Laurel A. Donnell-Fink, Naghmeh Dorrani, Daniel C. Dorset, Emilie D. Douine, David D. Draper, David J. Eckstein, Lisa T. Emrick, Christine M. Eng, Ascia Eskin, Cecilia Esteves, Tyra Estwick, Carlos Ferreira, Brent L. Fogel, Noah D. Friedman, William A. Gahl, Emily Glanton, Rena A. Godfrey, David B. Goldstein, Sarah E. Gould, Jean-Philippe F. Gourdine, Catherine A. Groden, Andrea L. Gropman, Melissa Haendel, Rizwan Hamid, Neil A. Hanchard, Lori H. Handley, Matthew R. Herzog, Ingrid A. Holm, Jason Hom, Ellen M. Howerton, Yong Huang, Howard J. Jacob, Mahim Jain, Yong-hui Jiang, Jean M. Johnston, Angela L. Jones, Isaac S. Kohane, Donna M. Krasnewich, Elizabeth L. Krieg, Joel B. Krier, Seema R. Lalani, C. Christopher Lau, Jozef Lazar, Brendan H. Lee, Hane Lee, Shawn E. Levy, Richard A. Lewis, Sharyn A. Lincoln, Allen Lipson, Sandra K. Loo, Joseph Loscalzo, Richard L. Maas, Ellen F. Macnamara, Calum A. MacRae, Valerie V. Maduro, Marta M. Majcherska, May Christine V. Malicdan, Laura A. Mamounas, Teri A. Manolio, Thomas C. Markello, Ronit Marom, Julian A. Martínez-Agosto, Shruti Marwaha, Thomas May, Allyn McConkie-Rosell, Colleen E. McCormack, Alexa T. McCray, Matthew Might, Paolo M. Moretti, Marie Morimoto, John J. Mulvihill, Jennifer L. Murphy, Donna M. Muzny, Michele E. Nehrebecky, Stan F. Nelson, J. Scott Newberry, John H. Newman, Sarah K. Nicholas, Donna Novacic, Jordan S. Orange, J. Carl Pallais, Christina G.S. Palmer, Jeanette C. Papp, Neil H. Parker, Loren D.M. Pena, John A. Phillips III, Jennifer E. Posey, John H. Postlethwait, Lorraine Potocki, Barbara N. Pusey, Chloe M. Reuter, Amy K. Robertson, Lance H. Rodan, Jill A. Rosenfeld, Jacinda B. Sampson, Susan L. Samson, Kelly Schoch, Molly C. Schroeder, Daryl A. Scott, Prashant Sharma, Vandana Shashi, Rebecca Signer, Edwin K. Silverman, Janet S. Sinsheimer, Kevin S. Smith, Rebecca C. Spillmann, Kimberly Splinter, Joan M. Stoler, Nicholas Stong, Jennifer A. Sullivan, David A. Sweetser, Cynthia J. Tifft, Camilo Toro, Alyssa A. Tran, Tiina K. Urv, Zaheer M. Valivullah, Eric Vilain, Tiphanie P. Vogel, Colleen E. Wahl, Nicole M. Walley, Chris A. Walsh, Patricia A. Ward, Katrina M. Waters, Monte Westerfield, Anastasia L. Wise, Lynne A. Wolfe, Elizabeth A. Worthey, Shinya Yamamoto, Yaping Yang, Guoyun Yu, Diane B. Zastrow, and Allison Zheng.

The Program for Undiagnosed Diseases (UD-PrOZA) co-investigators are Steven Callens, Paul Coucke, Bart Dermaut, Dimitri Hemelsoet, Bruce Poppe, Wouter Steyaert, Wim Terryn, and Rudy Van Coster.

## Acknowledgments

This work is funded by the Undiagnosed Diseases Network U01HG007672 to VS and DBG, U01HG007703 to SFN, and U54NS093793 to HJB, SY & MJW. HJB is also supported by R01GM067858 and R24OD022005, and SY & MFW by a Simons Foundation Functional Screen Award (368479). We would like to thank Alejandro Lomniczi (OHSU) for sharing the *IRF2BPL* cDNA and Hongling Pan for technical assistance. Research reported in this publication was supported by the National Institute of Neurological Disorders and Stroke (NINDS) under award number K08NS092898 and Jordan’s Guardian Angels (to G.M.). Confocal microscopy at BCM is supported in part by U54HD083092 to the Intellectual and Developmental Disabilities Research Center (IDDRC) Neurovisualization Core. HJB is an Investigator of the Howard Hughes Medical Institute. The content is solely the responsibility of the authors, and does not necessarily represent the official views of the National Institutes of Health. The funding sources had no role in the design and conduct of the study, collection, management, analysis and interpretation of the data, preparation, review or approval of the manuscript, or decision to submit the manuscript for publication. The authors would like to thank the subjects and their families for their participation.

## Web Resources

GeneMatcher, http://www.genematcher.org/

OMIM, http://www.omim.org/

Undiagnosed Diseases Network, https://undiagnosed.hms.harvard.edu

UDN *IRF2BPL* page, https://undiagnosed.hms.harvard.edu/genes/irf2bpl/

MARRVEL, http://www.marrvel.org/

ExAC, http://exac.broadinstitute.org/

gnomAD, http://gnomad.broadinstitute.org/

